# Epigenetic Reprogramming of Host Chromatin by the Transforming Parasite *Theileria annulata*

**DOI:** 10.1101/2025.10.27.684985

**Authors:** Sakshi Singh, Debabrata Dandasena, Akash Suresh, Madhusmita Subudhi, Sonam Kamble, Shweta Murthy, Vasundhra Bhandari, Paresh Sharma

**Affiliations:** National Institute of Animal Biotechnology, Hyderabad; Graduate Studies, Regional Centre for Biotechnology (RCB), Faridabad, India; National Institute of Pharmaceutical Education and Research (NIPER), Hyderabad

**Keywords:** Epigenetic Parasitism, Chromatin Remodeling, Host Cell Transformation, *Theileria annulata* Infection

## Abstract

*Theileria annulata*, a transforming apicomplexan parasite, extensively reprograms the chromatin architecture of bovine leukocytes to facilitate infection and cellular transformation. To elucidate the underlying epigenetic mechanisms, we characterized the chromatin landscape of infected lymphocytes using integrated proteomic, imaging, and functional assays. Our findings reveal that *T. annulata* lacks the DNA damage marker γH2A.X and its associated SQ/TQ motif, indicating an evolutionary divergence from canonical DNA repair signalling pathways.

High-resolution profiling of histone post-translational modifications (PTMs) demonstrated distinct nuclear compartmentalization, with host (H3K27me3, H3K9me1/2), parasite-predominant modifications (H3K4me3, H3K18me1, H3K27ac, H3K9ac, and H3K36me3), and shared modifications (e.g., H3K4me1/2, H3K36me2, H4K5/8/12/16ac, H3K9me3 and H4K91Ac) suggesting a coordinated epigenetic strategy driving host cell programs to sustain proliferation and survival. Our study delineates a novel host–parasite epigenetic interface, positioning *T. annulata* as a unique model of epigenetic parasitism. By bridging parasitology and cancer epigenetics, these findings unveil new therapeutic opportunities targeting chromatin vulnerabilities in parasite-induced transformation.

**Graphical abstract:** 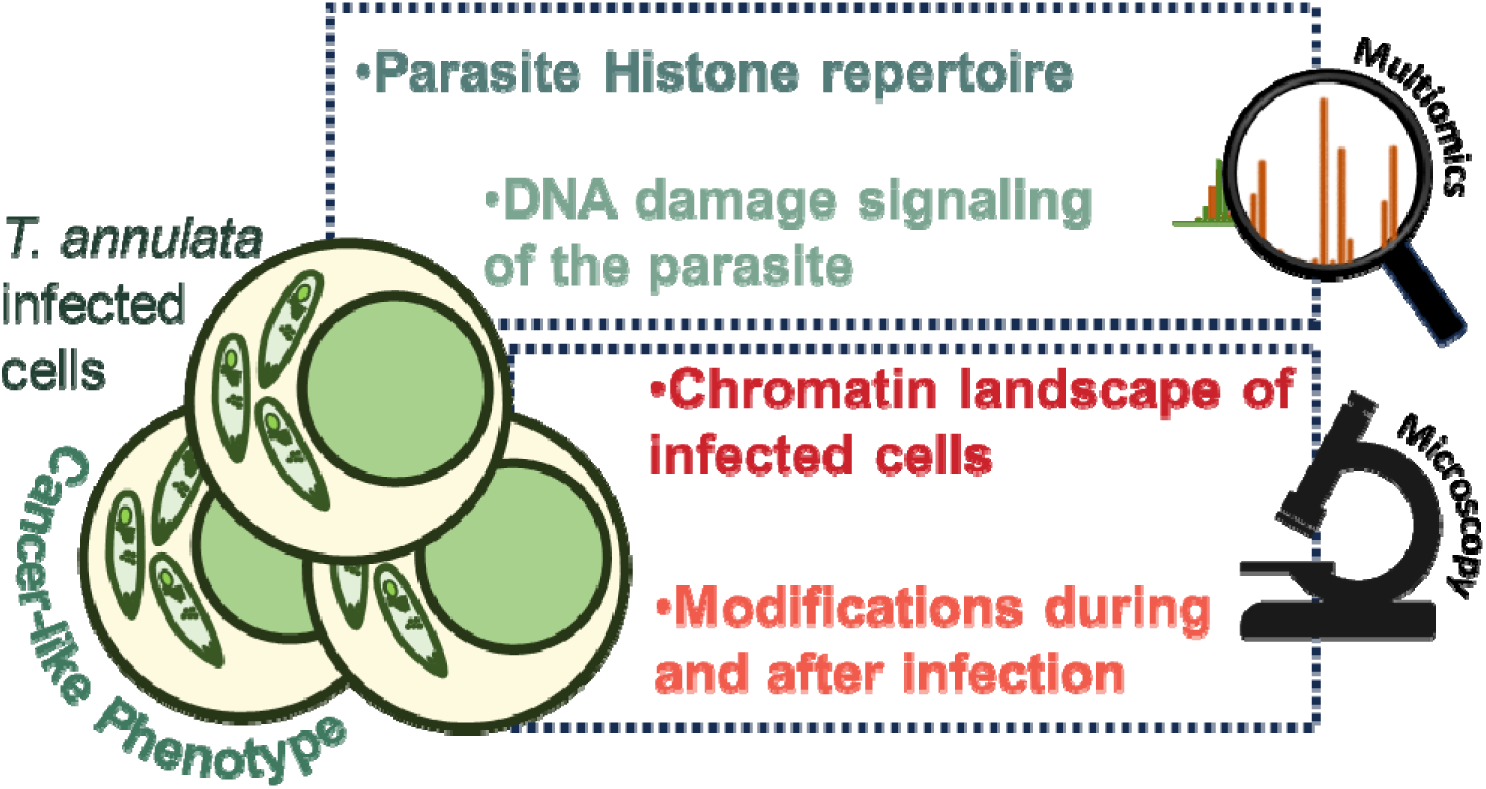

## Introduction

*Theileria annulata* is a tick-transmitted apicomplexan parasite and the etiological agent of bovine theileriosis (BT), a disease that imposes substantial economic burdens on livestock industries across Africa, Asia, and the Middle East (Kumar et al., 2017). In India alone, BT accounts for over $1.2 billion in annual losses due to decreased productivity, elevated mortality rates, and treatment-related expenses (Narladkar et al., 2018).

Unlike most intracellular pathogens, *T. annulata* possesses a unique capacity to induce a reversible, cancer-like transformation of infected bovine leukocytes (Tretina et al., 2015). These transformed cells exhibit hallmark features of malignancy, including uncontrolled proliferation, resistance to apoptosis, and invasive behavior (Woods et al., 2021). The emergence of *T. annulata* strains resistant to buparvaquone (BPQ), the frontline anti-theilerial drug, underscores the urgent need for alternative therapeutic strategies (Mhadhbi et al., 2010).

Previous studies have demonstrated that *T. annulata* activates host oncogenic signaling pathways such as *PI3K-AKT, NF-*_κ_*B, ERK1/2, and JAK-STAT, while concurrently suppressing* the tumor suppressor *TP53* (Heussler et al., 2001; Dessauge et al., 2005; Haller et al., 2010; Barman et al., 2023; Dandesena et al., 2025). However, the epigenetic mechanisms underpinning this transformation remain inadequately characterized. Given the striking parallels between parasite-induced transformation and tumorigenesis, it is plausible that *T. annulata* exploits host chromatin regulators to reprogram gene expression and maintain its intracellular niche.

Histone variants, post-translational modifications (PTMs)—including acetylation, methylation, and phosphorylation—and their associated modifying enzymes are now recognized as central regulators of chromatin architecture, transcriptional control, and cell fate decisions (Bannister & Kouzarides, 2011; Zhao & Shilatifard, 2019). In oncology, targeting chromatin modifiers has emerged as a viable strategy to reverse malignant epigenetic states (Ghosh et al., 2021; Yu et al., 2024).

Similarly, the expanding field of patho-epigenetics reveals how pathogens leverage epigenetic plasticity to modulate virulence, antigenic variation, and persistence (Schator et al., 2021). In apicomplexan parasites such as *Plasmodium falciparum* and *Toxoplasma gondii*, chromatin remodeling is essential for developmental transitions and immune evasion (Croken et al., 2012). Despite these advances, the extent to which *T. annulata* manipulates both host and parasite chromatin landscapes to drive cellular transformation remains largely unexplored.

This study addresses this critical gap by systematically profiling the histone composition and PTM landscape in *T. annulata*-infected cells (TA cells). We investigate the epigenetic interface between host and parasite, elucidating its impact on key regulatory pathways, including apoptosis. Our findings not only enhance the mechanistic understanding of *T. annulata* pathogenesis but also offer broader insights into how intracellular pathogens hijack host epigenetic machinery to promote survival and sustain a cancer-like phenotype.

## Results

### 1. Uncovering the Epigenetic Blueprint of *T. annulata*: Histone Variants and Their Evolutionary Trajectories

To elucidate how *T. annulata* modulates host chromatin and gene expression, we first characterized the parasite’s histone repertoire—an essential step given the limited annotation of histone proteins in this organism. Histones were isolated from TA cells (Fig. 1a) and analyzed via LC-MS/MS (Fig. 1b). Peptide mapping against host and parasite proteomes identified seven parasite-derived histones, including canonical core histones (H2A, H2B, H3, H4) and variants (H2A.Z, H2B.Z, H3.3).

**Figure 1.**
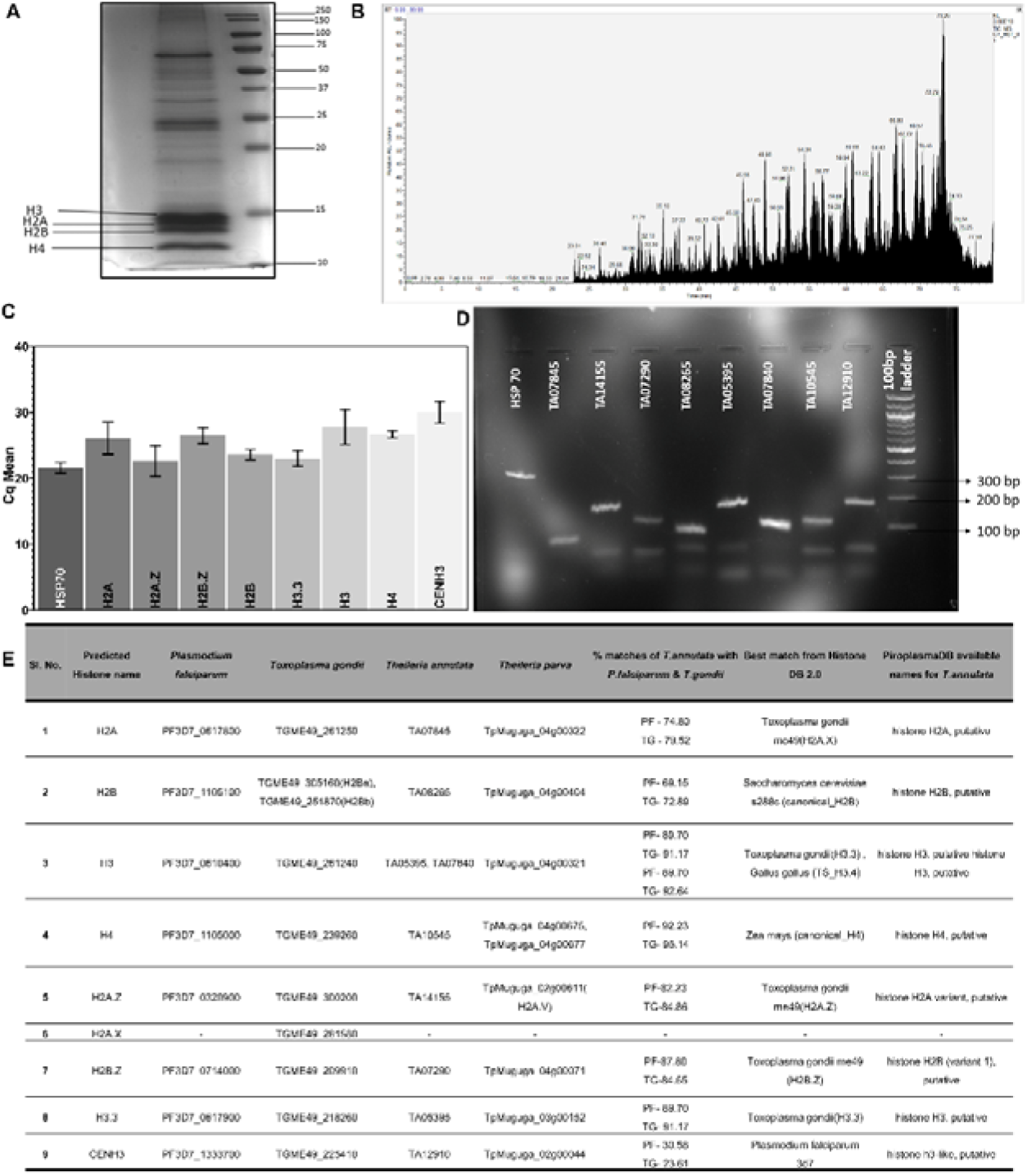
Molecular Characterization and Evolutionary Analysis of *T. annulata* Histones. A. Coomassie-stained SDS-PAGE gel demonstrating successful isolation of histone proteins from TA cells. B. Representative chromatogram from mass spectrometry analysis of purified histone proteins. C. Bar graph showing mean quantification cycle (Cq) values for candidate histone genes (TA07845, TA14155, TA07290, TA08265, TA05395, TA07840, TA10645, TA12310) alongside the parasite reference gene HSP70. Error bars indicate standard deviation from n = 2 independent biological replicates. D. Agarose gel electrophoresis (1.8%) of qPCR products stained with ethidium bromide and visualized under UV illumination. Bands at expected sizes confirm successful amplification and transcription of target genes. E. Summary table of predicted histone variants in *T. annulata*, identified via sequence comparisons with histone genes from *P. falciparum, T. gondii*, and *T. parva*. BLAST searches and annotations from PiroplasmaDB, PlasmoDB, and ToxoDB were used. HistoneDB 2.0 was employed to determine the best-matched organism and canonical or variant classification. For each predicted histone, the closest ortholog, percent identity, and putative annotation are listed.

To confirm the parasite origin and exclude host contamination, expression of *T. annulata* histone genes was validated by qPCR (Fig. 1c–d). No expression was detected in uninfected bovine PBMCs (bPBMCs), supporting their parasite-specific origin.

Comparative BLAST analysis against *Plasmodium* and *Toxoplasma* histone databases confirmed the presence of conserved histones and variants (Fig. 1e). While most histones exhibited high sequence conservation with other apicomplexans, two notable differences emerged: (1) absence of H2A.X, a variant critical for DNA damage signaling, and (2) low sequence similarity (23–30%) of CEN H3 with orthologs from related species. No linker histone H1 was detected, consistent with prior findings in *Plasmodium* (Hernandez-Rivas et al., 2010).

Further analysis using Histone Database 2.0 revealed that while most *T. annulata* histones closely resembled those of *T. gondii*, H2B aligned more closely with *Saccharomyces cerevisiae*, H4 with *Zea mays*, and CEN H3 with *P. falciparum* (Fig. 1e), highlighting both evolutionary conservation and divergence in the parasite’s chromatin machinery.

LC-MS/MS-based PTM analysis revealed a diverse array of modifications in parasite histones, including acetylation, methylation, ubiquitination, oxidation, crotonylation, propionylation, and formylation. In contrast, host histones exhibited additional modifications such as phosphorylation, succinylation, and sulfation (Supplementary Table 1), suggesting a distinct regulatory environment.

### 2. Absence of H2A.X and Loss of Canonical DNA Damage Signaling in *T. annulata*

Given the central role of γH2A.X in DNA damage response across eukaryotes (Kinner et al., 2008), we investigated whether *T. annulata* encodes this histone variant. Immunofluorescence assays (IFA) using a human γH2A.X antibody revealed strong nuclear staining in host cells but no detectable signal in the parasite, even following etoposide-induced DNA damage (Fig. 2a). This antibody is known to cross-react with H2A.X in *Toxoplasma* (Velásquez et al., 2024), H2A in *Plasmodium* (Goyal et al., 2021), and bovine H2A.X, suggesting either absence or extreme divergence in *T. annulata*.

**Figure 2.**
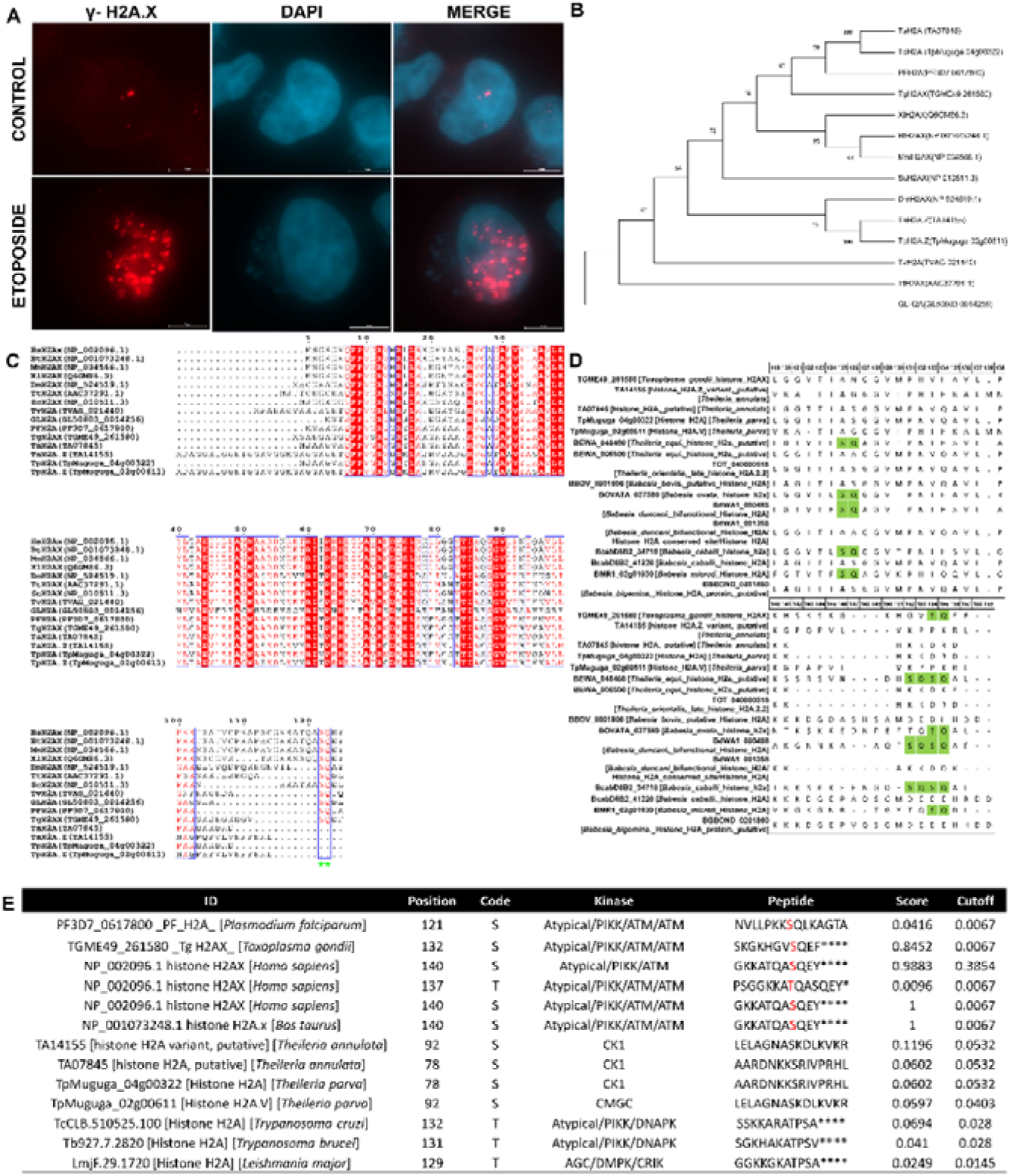
Evolutionary Conservation and Functional Analysis of _γ_H2A.X in *T. annulata*. A. Immunofluorescence assay (IFA) showing γH2A.X expression in TA control cells and cells treated with etoposide (10 µM, 24 hr). B. Maximum likelihood phylogenetic tree depicting evolutionary relationships of *T. annulata* histone H2A (TA07845) with H2A and H2A.X orthologs from protozoan parasites and selected eukaryotes. Organisms and gene IDs include: *T. annulata* (TA07845, TA14155), *T. parva* (TpMuguga_04g00322, TpMuguga_02g00611), *P. falciparum* (PF3D7_0617800), *T. gondii* (TGME49_261580), Homo sapiens (NP_002096.1), *Bos taurus* (NP_001073248.1), *S. cerevisiae* (NP_010511.3), *D. melanogaster* (NP_524519.1), *T. thermophila* (AAC37291.1), *T. vaginalis* (TVAG_021440), *G. lamblia* (GL50803_0014256), *X. laevis* (Q6GM86.3), *M. musculus* (NP_034566.1). C. Multiple sequence alignment of H2A.X variants using Clustal Omega, visualized with ESPript 3.0. Conserved regions are highlighted, with emphasis on SQ motifs (boxed in blue and marked with green stars), which are potential ATM/ATR phosphorylation sites. The canonical γH2A.X motif (SQE/DΦ) is conserved in several protozoan orthologs. Fully conserved residues are shown in white on red; similar residues are shaded in red. D. Alignment of H2A sequences from apicomplexan parasites to assess conservation of the serine-glutamine (SQ) motif—a hallmark of H2A.X and a known target of ATM/ATR kinases. Species analyzed include *T. gondii*, *T. annulata*, *T. parva*, *T. equi*, *T. orientalis*, *B. bovis*, *B. ovata*, *B. duncani*, *B. caballi*, *B. microti*, and *B. bigemina*. Conserved residues are indicated; SQ motifs are highlighted in green. This analysis reveals species-specific presence or absence of γH2A.X-like domains, suggesting evolutionary divergence in DNA damage signaling across Apicomplexa. E. Predicted phosphorylation sites on H2A.X and related variants using GPS 6.0. Shown are kinase–substrate interactions, peptide sequences, phosphorylation positions, kinase types, and confidence scores. Sequences analyzed include canonical and variant H2A from *P. falciparum, T. gondii, H. sapiens, B. taurus, T. annulata, T. parva, T. cruzi, T. brucei,* and *L. major*.

Phylogenetic analysis of *T. annulata* H2A (TA07845) revealed clustering with the H2A.X variant of *T. gondii* and canonical H2A of *P. falciparum*, but not with H2A.X from higher eukaryotes (Fig. 2b). Multiple sequence alignment (Fig. 2c) showed that the conserved C-terminal SQ motif—required for ATM/ATR-mediated phosphorylation and γH2A.X formation—is absent in *T. annulata* (Marechal & Zou, 2013). A similar pattern was observed in *Theileria parva*, suggesting a lineage-specific loss.

To determine whether this absence is a broader feature of piroplasms, we extended our analysis to additional species. While *Theileria equi, Babesia ovata, Babesia duncani, Babesia caballi,* and *Babesia microti* retained the SQ motif, *T. annulata, T. parva, Theileria orientalis, Babesia bovis,* and *Babesia bigemina* lacked it entirely (Fig. 2d).

To explore potential compensatory mechanisms, GPS 6.0 was used to predict serine/threonine/tyrosine phosphorylation sites in *T. annulata* H2A. Although two serine residues (S78 and S92) were predicted to be phosphorylated, they were located outside the C-terminal region and associated with CK1 and CMGC kinase families (Fig. 2e). These kinases have been implicated in DNA damage responses in other systems (Cullati et al., 2023; Kciuk et al., 2022), but the low prediction scores suggest limited functional relevance.

Additionally, we examined DNA damage markers in *Trypanosoma cruzi, Trypanosoma brucei,* and *Leishmania major*, where threonine phosphorylation has been proposed as a marker of DNA damage. Although threonine and tyrosine phosphorylation sites were identified, they lacked positional relevance and kinase specificity.

Collectively, these findings indicate that *T. annulata, T. parva,* and *T. orientalis* lack the canonical γH2A.X-mediated DNA damage signaling pathway. The absence of the SQ/TQ motif and lack of ATM/ATR-targeted phosphorylation sites support the hypothesis that these parasites may have evolved an alternative, yet uncharacterized mechanism to detect and respond to genomic stress.

### 3. Differential Distribution of Histone Modifications in Host and Parasite Chromatin

To investigate the epigenetic landscape of TA cells, we analyzed the localization of key histone PTMs using IFA with a panel of 18 well-characterized antibodies targeting lysine acetylation (n=5) and methylation marks (n=13) on H3 and H4. These modifications were selected based on their established roles in gene regulation in both apicomplexan parasites and cancer (Supplementary Table 1). Although most antibodies were raised against human histones, they exhibited strong cross-reactivity with both host and parasite, enabling comparative analysis.

PTMs were categorized based on nuclear distribution patterns: (1) predominantly host-enriched, (2) evenly distributed between host and parasite, or (3) parasite-enriched (Fig. 3-6). Repressive marks such as H3K9me2, H3K9me1, and H3K27me3 showed significantly higher intensity in host nuclei, while the activation mark H3K79me2 was also host-biased (Fig. 3a). Seven modifications—including activation-associated marks (H3K4me1, H3K36me2, H4K5/8/12/16ac, H4K16ac, H3K4me2, and H4K91ac) and the repressive mark H3K9me3— displayed balanced distribution across both nuclei (Fig. 4).

**Figure 3.**
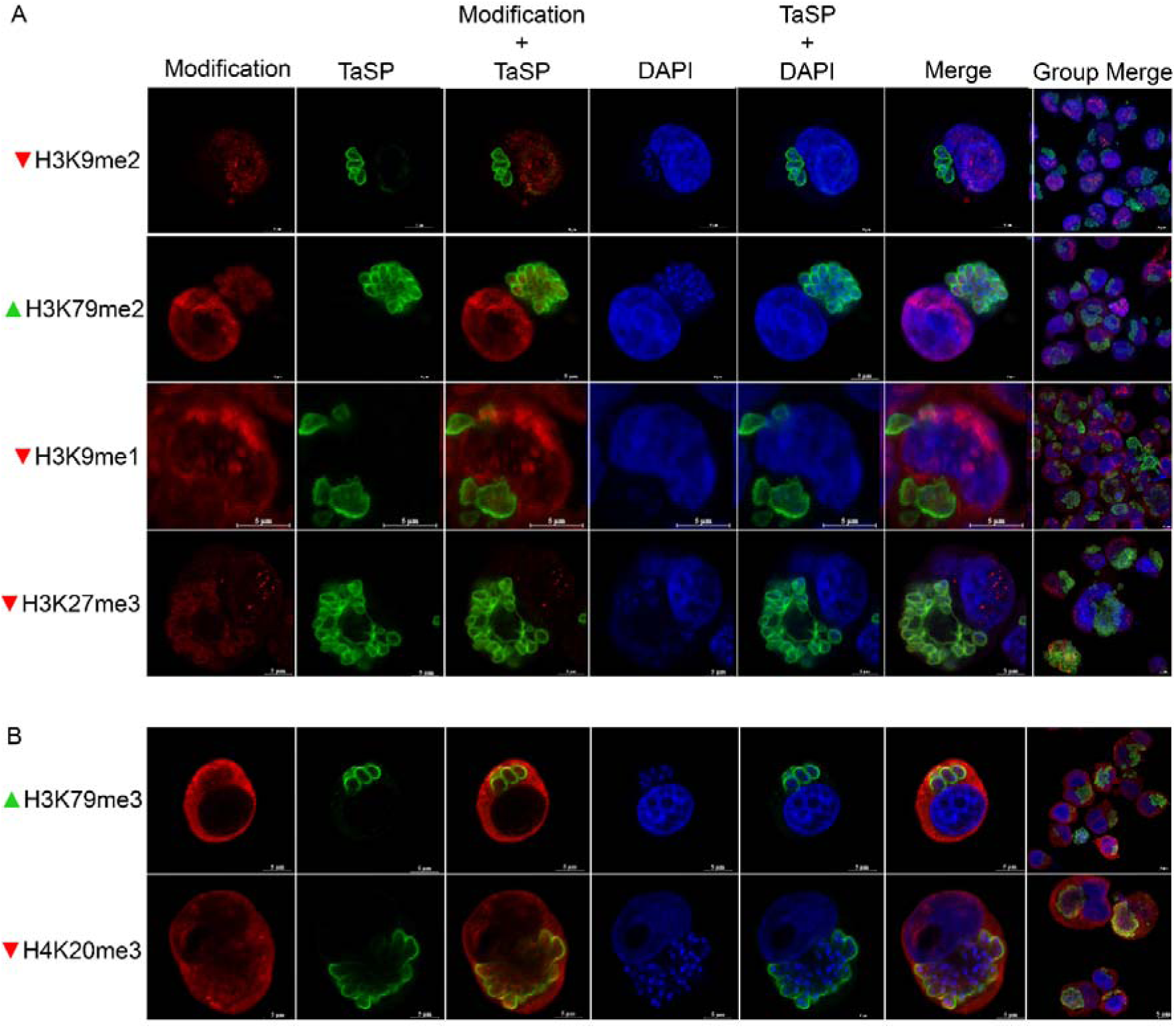
Subcellular Localization of Histone Methylation Marks in TA Cells IFA showing nuclear expression of histone methylation marks in host cells: H3K9me2, H3K79me2, H3K9me1, and H3K27me2 (Panel A). Cytoplasmic localization of H3K79me3 and H4K20me3 is shown in Panel B.

**Figure 4.**
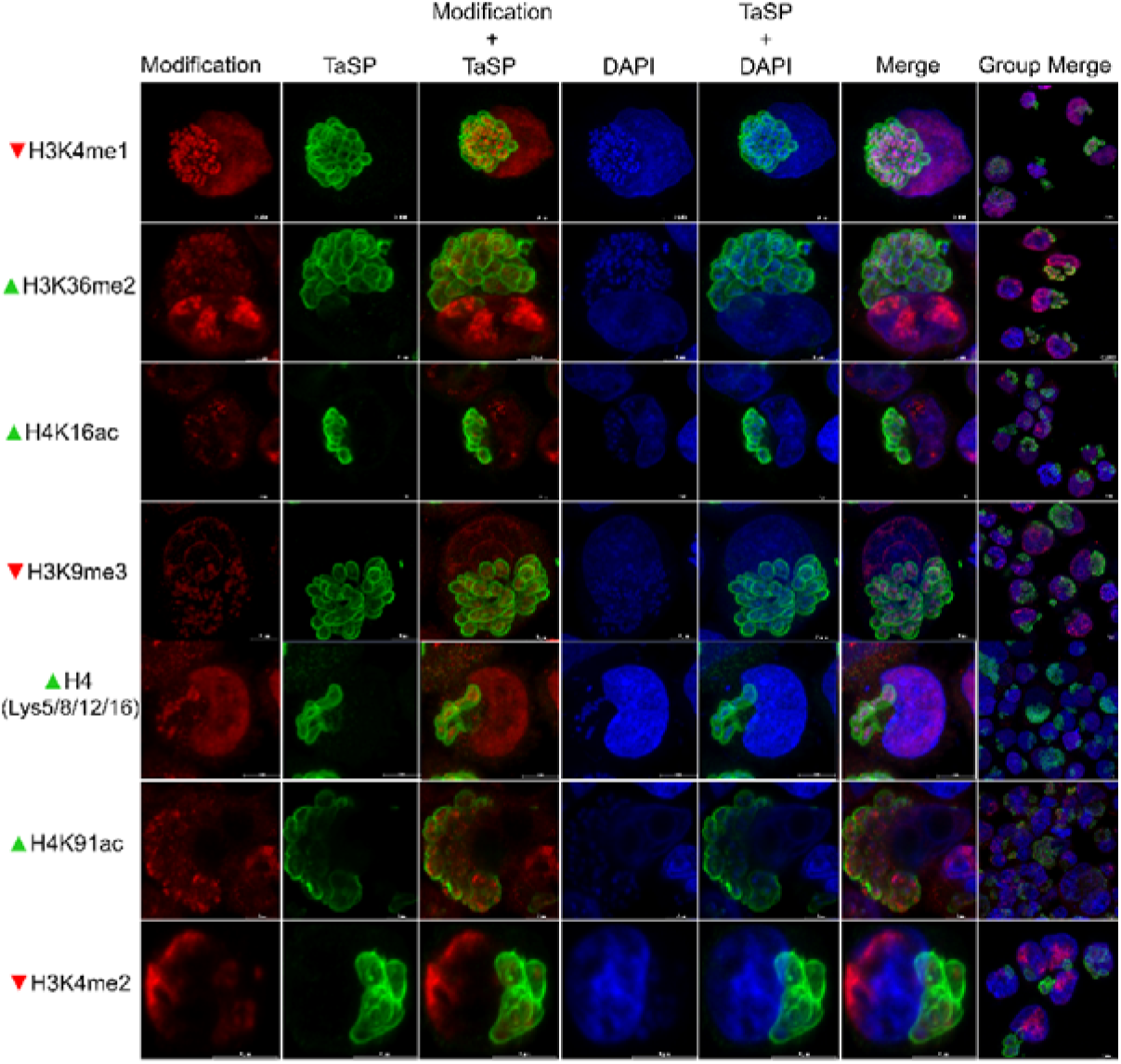
IFA of Host and Parasite-Associated Histone Modifications in TA Cells IFA displaying histone modifications with shared localization in host and parasite nuclei: H3K4me1, H3K36me2, H4K16ac, H3K9me3, H4K5/8/12/16ac, H4K91ac, and H3K4me2.

Five histone marks were significantly enriched within the parasite: activation-associated H3K4me3, H3K27ac, H3K9ac, and H3K36me3, along with the repressive mark H3K18me1. To determine whether these PTMs were parasite-driven, TA cells were treated with BPQ, which eliminates *T. annulata*. IFA revealed a dramatic reduction in all parasite-enriched marks (Fig. 5-6).

**Figure 5.**
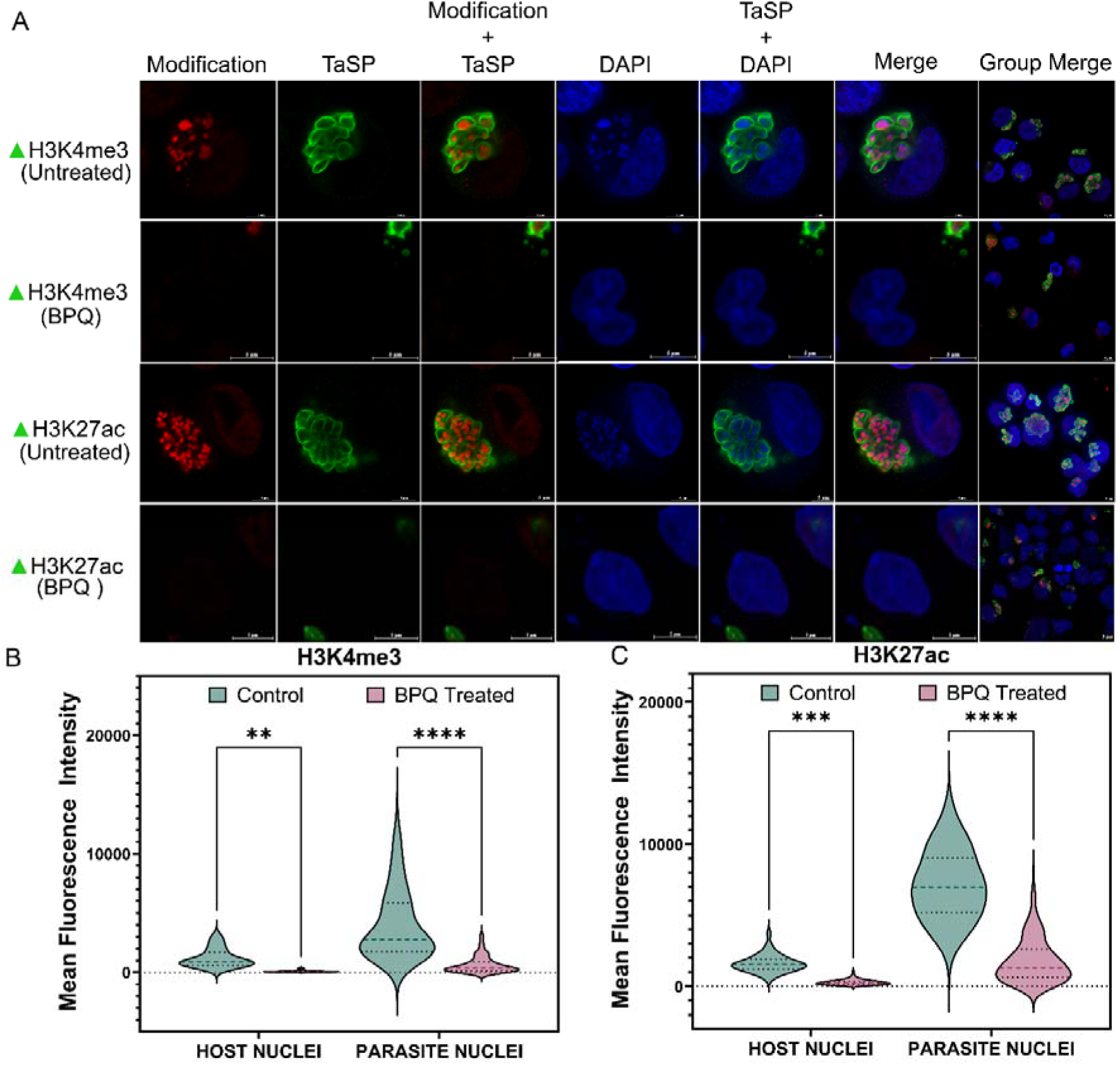
Quantitative Analysis of H3K4me3 and H3K27ac in TA Cells. A. IFA showing expression of H3K4me3 and H3K27ac in TA control cells and cells treated with buparvaquone (BPQ; 200 ng/mL, 96 hr). B. Quantification of H3K4me3 expression in host and parasite nuclei: control host (n = 77), control parasite (n = 146); BPQ-treated host (n = 46), BPQ-treated parasite (n = 118). C. Quantification of H3K27ac expression: control host (n = 40), BPQ-treated host (n = 99); control parasite (n = 197), BPQ-treated parasite (n = 230). Statistical analysis was performed using two-way ANOVA. Significance is indicated as follows: **P = 0.0024; ***P = 0.0001; ****P < 0.0001.

**Figure 6.**
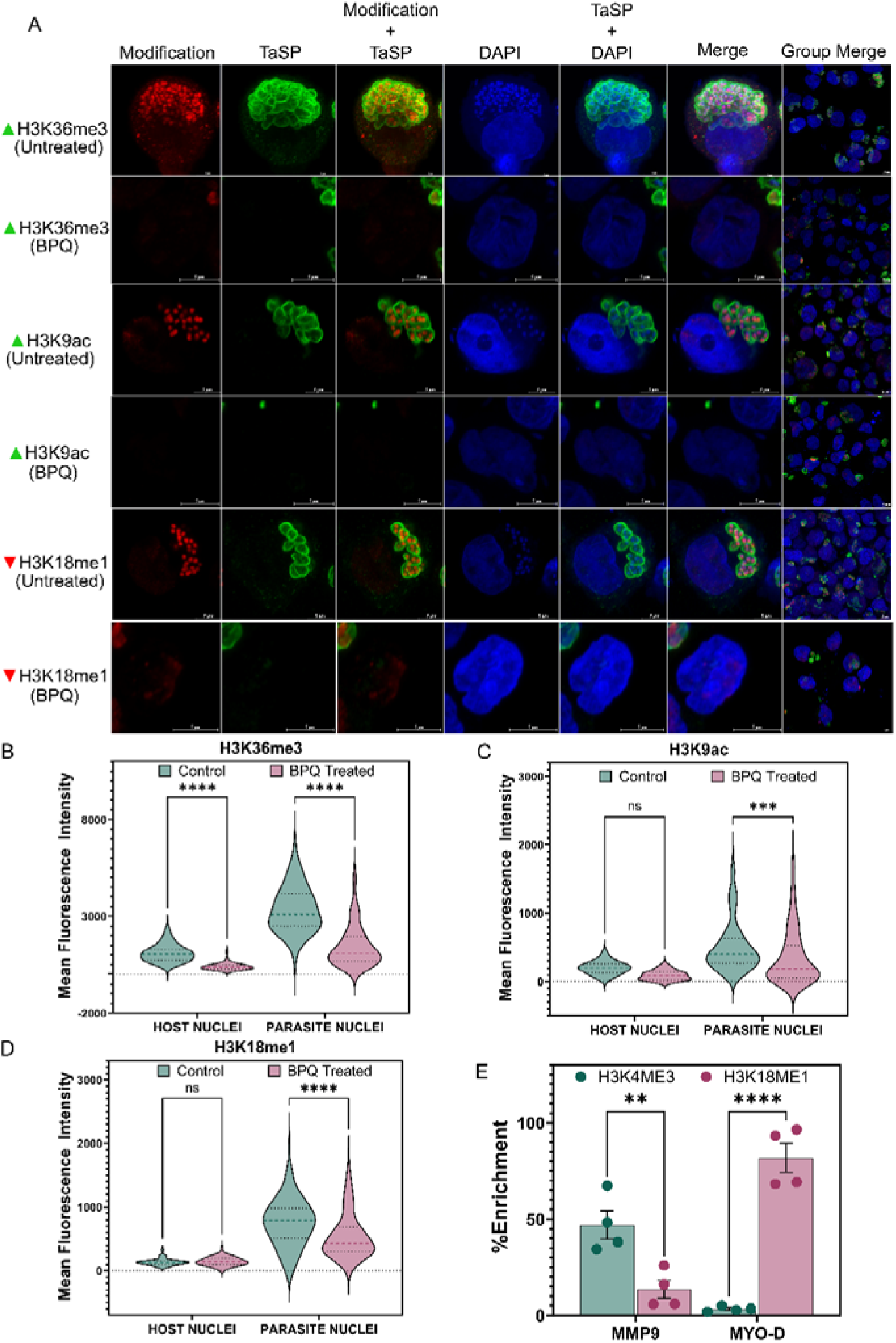
Impact of BPQ Treatment on Parasite-Associated Histone Modifications and Enrichment on Host Genes in TA Cells. A. Immunofluorescence assay (IFA) showing expression of histone marks H3K36me3, H3K9ac, and H3K18me1 in TA control and BPQ-treated cells (200 ng/mL, 96 hr). B–D. Quantitative analysis of histone mark intensity in host and parasite nuclei: B. H3K36me3 (control host nuclei, n = 54; BPQ host nuclei, n = 77; control parasite nuclei, n = 113; BPQ parasite nuclei, n = 160). C. H3K9ac (control host nuclei, n = 22; BPQ host nuclei, n = 55; control parasite nuclei, n = 188; BPQ parasite nuclei, n = 162). D. H3K18me1 (control host nuclei, n = 48; BPQ host nuclei, n = 62; control parasite nuclei, n = 104; BPQ parasite nuclei, n = 145). In the graphs: ns indicates P = 0.6997; *** indicates P = 0.0088; **** indicates P < 0.0001. E. Bar graph showing percentage enrichment of H3K4me3 and H3K18me1 at host gene promoters MMP9 and MYO-D. Two-way ANOVA indicates significant differences: **P = 0.0031; ****P < 0.0001.

To further validate their origin, we examined PTM levels in uninfected controls. The abundance of these marks was markedly lower in uninfected cells, indicating that the majority of the detected modifications originated from the parasite (Supplementary Fig. 1). This confirms that *T. annulata* actively contributes to histone modification, potentially to manipulate host gene expression. Parasite clearance led to cell death, suggesting functional relevance.

H3K79me3 and H4K20me3 localized exclusively to the cytoplasm, suggesting lack of nuclear binding or antibody accessibility (Fig. 3b).

**Supplementary Figure 1:**
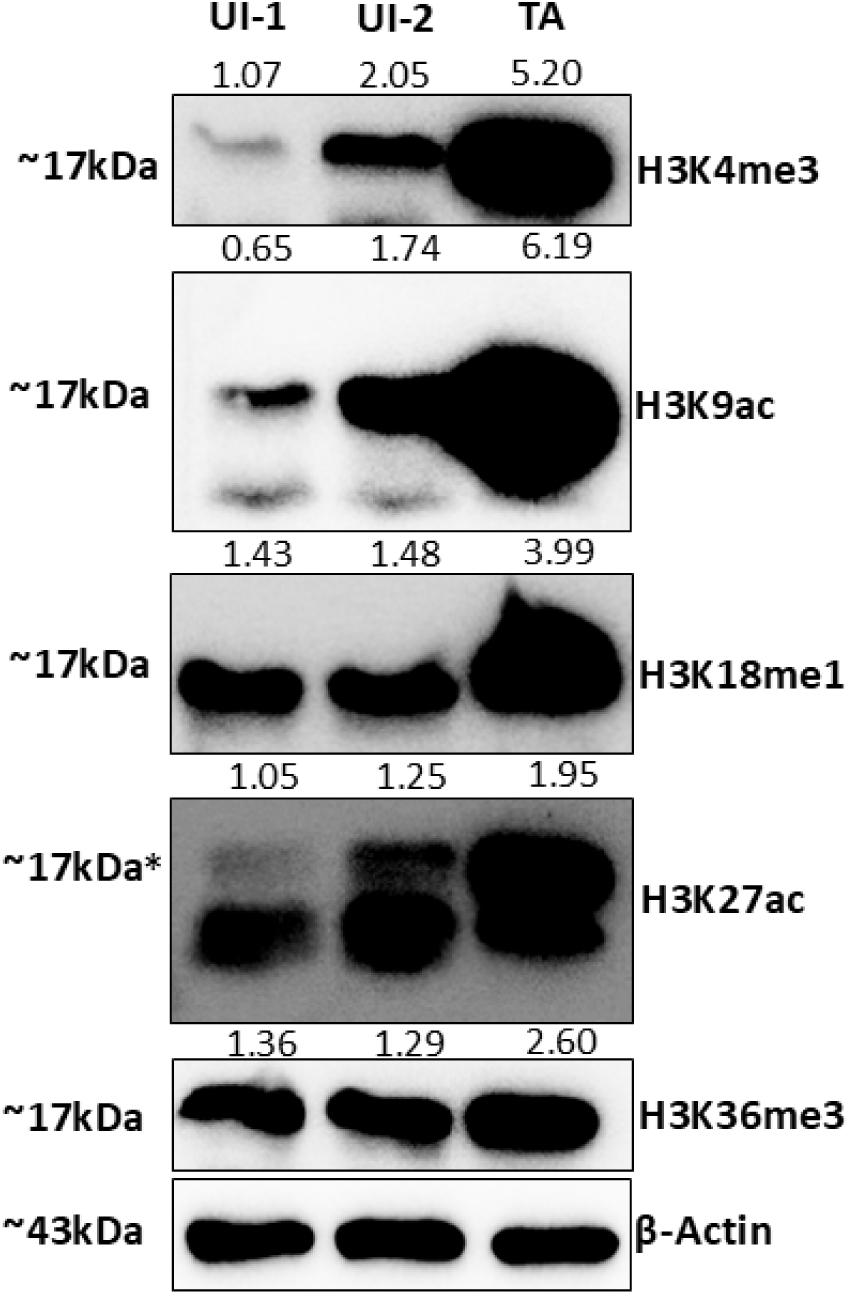
Difference in the expression levels of parasite-predominant marks between uninfected and infected cells. Western blot analysis comparing parasite-predominant histone marks in uninfected (n = 2) versus infected cells. * denotes right band size.

### 4. Differential Localization of Histone Marks Highlights Model-Specific Epigenetic Variability

Previous work by Cheeseman et al. (2021) characterized the distribution of histone marks H3K4me3, H3K36me3, and H3K18me1 in TA cells, reporting uniform nuclear localization of H3K4me3 and H3K36me3 across host and parasite nuclei, and parasite-specific enrichment of H3K18me1. In contrast, our IFA analysis revealed a distinct pattern: all three marks were predominantly enriched within the parasite nucleus, with only low but detectable levels present in host nuclei. These discrepancies may reflect differences in infection models or antibody specificity.

To validate our observations, we performed ChIP-qPCR at the promoter of the host gene MYO-D, which showed enrichment for the repressive mark H3K18me1, consistent with our IFA findings. As a positive control, we examined the MMP9 promoter, a metastasis-associated gene previously linked to H3K4me3 in TA cells (Cock-Rada et al., 2012). As expected, H3K4me3 was enriched at this locus, while H3K18me1 was depleted (Fig. 6e). These results suggest model-specific epigenetic variability: while Cheeseman et al. utilized *in vitro*-infected cell lines, our study employed established cell lines derived from field-isolated clinical samples.

## Discussion

This study uncovers a multifaceted epigenetic strategy by which *T. annulata* reshapes host chromatin to sustain cellular transformation. Our integrative analysis highlights a distinct host–parasite interface governed by chromatin remodeling, providing mechanistic insights into parasite-induced immortality.

A striking observation is the absence of H2A.X and its SQ/TQ motif in *T. annulata*, a key component of the DNA damage response in eukaryotes (Podhorecka et al., 2010; Marechal & Zou, 2013). Unlike related apicomplexans such as *P. falciparum* and *T. gondii*, which retain this variant (Dalmasso et al., 2009; Goyal et al., 2021), *T. annulata* appears to have diverged evolutionarily, potentially enabling evasion of host genomic surveillance. However, the mechanisms by which the parasite maintains its own genome integrity remain to be elucidated.

Histone post-translational modifications (PTMs) play a pivotal role in regulating gene expression. In TA cells, we analyzed 18 distinct histone marks, categorizing them based on their localization in the host nucleus, parasite nucleus, or shared between both compartments. The presence of shared histone modifications (e.g., H3K4me1, H3K36me2, H4K16ac) suggests the existence of a cross-species regulatory network enabling coordinated transcriptional control. Additionally, the coexistence of activating and repressive marks within host chromatin (e.g., H3K79me2, H3K9me1/2, H3K27me3) indicates a dynamic balance between sustaining essential host functions and restricting immune responses. These findings highlight chromatin plasticity as a therapeutic vulnerability in parasite-induced transformation.

The chromatin landscape of *T. annulata* is characterized by high levels of transcriptionally active histone marks such as H3K4me3, H3K27ac, H3K36me3, and H3K9ac, along with the repressive mark H3K18me1. This pattern aligns with chromatin strategies observed in other apicomplexans (Nardelli et al., 2013; Serrano-Durán et al., 2022), suggesting that *Theileria* may mirror mechanisms used by *Plasmodium* and *Toxoplasma* to regulate virulence and immune evasion (Salcedo-Amaya et al., 2009; Ukaegbu et al., 2014; Duffy et al., 2017; Pickford et al., 2021). Similar chromatin remodeling has also been documented in viral oncogenesis, where pathogens restructure host nuclear architecture to modulate gene expression (Rehman et al., 2023).

Despite these advances, the study has limitations. The reliance on antibody-based detection may be affected by cross-reactivity, and the lack of parasite-specific ChIP-grade reagents limits resolution. Future studies should employ ChIP-seq and proteomic approaches to map chromatin regulators more precisely. Identifying parasite effectors that initiate host chromatin remodeling will be critical to understanding the upstream signals driving these changes. Moreover, exploring similar epigenetic mechanisms in other intracellular pathogens could broaden our understanding of host–pathogen interactions.

## Conclusion

Our study establishes *T. annulata* as a potent epigenetic regulator that remodels host chromatin to inhibit apoptosis and drive a cancer-like phenotype. The absence of H2A.X, a key mediator of DNA damage signaling, suggests a unique genomic adaptation that may facilitate parasite persistence. TA cells exhibit a diverse array of histone modifications, potentially contributing to their transformed state. These findings advance our understanding of *T. annulata* pathogenesis and underscore the relevance of chromatin modifiers as viable drug targets. Future research should aim to characterize the molecular effectors driving these alterations and explore whether similar epigenetic mechanisms operate in other intracellular pathogens capable of inducing transformation.

## Materials and Methods

### Cell Culture and Drug Treatment

Peripheral blood mononuclear cells (PBMCs) were isolated from cattle exhibiting clinical symptoms of *T. annulata* infection. Following standard density gradient centrifugation, PBMCs were cultured in RPMI 1640 medium supplemented with 10% fetal bovine serum (FBS) and 100 µg/mL penicillin-streptomycin. Cultures were maintained at 37°C in a humidified incubator with 5% COC.

To confirm in vitro infection, PCR was performed using primers targeting the parasite-specific TaSP gene (Dandasena et al., 2018). Positive amplification validated the presence of *T. annulata* in cultured cells (TA cells).

As controls, PBMCs were also obtained from clinically healthy cattle (bPBMCs; n = 3), confirmed to be free of *T. annulata* infection. For treatment assays, TA cells were incubated with BPQ (200 ng/mL) for 96 hours and with Etoposide (10 µM) for 3 hours.

### Histone Extraction and Mass Spectrometry Analysis

Histones were extracted from TA cells (2 × 10C) using the EpiQuik™ Total Histone Extraction Kit (EpiGentek, OP-0006) according to the manufacturer’s protocol. Protein integrity was assessed by SDS-PAGE followed by Coomassie staining.

Fifty micrograms of histone extract were reduced with DL-dithiothreitol (Sigma, D9779) at 56 °C, alkylated with iodoacetamide (Sigma, I1149) at room temperature for 1 hour, and digested overnight with sequencing-grade trypsin (Promega, V5111) at 37 °C. The reaction was quenched with formic acid and centrifuged at 14,000 rpm for 10 minutes. Peptides were transferred to fresh tubes, dried using a SpeedVac, and desalted using C18 spin tips (Pierce, 87782).

Samples were analyzed on an Orbitrap Fusion mass spectrometer (Thermo Fisher Scientific). Data were processed using Proteome Discoverer 2.4 SP1 and searched against UniProt databases for *Bos taurus* (Proteome ID: UP000009136) and *T. annulata* (Proteome ID:

UP000001950). Search parameters included a 10 ppm precursor mass tolerance, allowance for two missed trypsin cleavages, and a false discovery rate (FDR) of 0.01–0.05.

Variable modifications included acetylation (lysine and arginine), mono-, di-, and tri-methylation (lysine and arginine), phosphorylation (serine, threonine, tyrosine), O-GlcNAc modification (serine/threonine), succinylation, propionylation, crotonylation, formylation, sulfation, ubiquitination (lysine), oxidation (methionine), and N-terminal acetylation.

### RNA Isolation and Quantitative PCR

Total RNA was extracted from TA cells and bPBMCs (5 × 10C) using the Macherey-Nagel RNA extraction kit (740984.250). cDNA synthesis was performed using the PrimeScript cDNA Synthesis Kit (Takara, 6110A). Gene expression was quantified using SYBR Green-based qPCR on a Bio-Rad CFX96 system.

Primer sequences are listed in Supplementary Table 1. HSP70 (parasite) and HPRT (host) were used as reference genes. PCR conditions included an initial denaturation at 95°C for 3 minutes, followed by 40 cycles of 95°C for 10 seconds and 55°C for 30 seconds, with melt curve analysis. Experiments were performed using n = 2 biological replicates.

### Bioinformatics

Histone sequences from *T. annulata, T. parva, P. falciparum, and T. gondii* were retrieved from PiroplasmDB, PlasmoDB, and ToxoDB. Sequences from *T. annulata* were also compared against entries in HistoneDB 2.0 (https://www.ncbi.nlm.nih.gov/research/HistoneDB2.0), a curated database of histone protein sequences across species.

Alignments were performed using ClustalW and visualized with ESPript3. Phylogenetic trees were constructed using MEGA4 (Tamura et al., 2007). Phosphorylation sites were predicted using GPS 6.0 (Xue et al., 2008), and conserved motifs were analyzed using Jalview.

### Immunofluorescence Assay (IFA) and Western Blotting

TA cells (5 × 10C), untreated or treated, were fixed with 4% paraformaldehyde and mounted on positively charged microscope slides. Cells were permeabilized with 0.1% Triton X-100 and blocked using a solution containing 1% BSA, 10% normal goat serum, and 0.3 M glycine in 0.1% PBS-Tween.

Cells were incubated with primary antibodies (Supplementary Table 1), followed by secondary antibodies and DAPI staining. Imaging was performed using a Carl Zeiss Airyscan fluorescence microscope, and fluorescence intensity was quantified using ZEN 3.3 software.

Western blotting was conducted using protein lysates (20–40 µg) from bPBMCs and TA cells. Samples were resolved by SDS-PAGE and transferred onto PVDF membranes. Membranes were blocked with 5% non-fat milk in TBST and incubated with primary antibodies (Supplementary Table 1). β-Actin was used as a loading control. Signal detection was performed using the Bio-Rad ChemiDoc™ imaging system.

### Chromatin Immunoprecipitation (ChIP)

ChIP was performed on TA cells (1 × 10C) fixed with 1% formaldehyde for 10 minutes at room temperature and quenched with 125 mM glycine. Cell lysis was carried out using Hi-C cell lysis buffer (10 mM Tris-Cl pH 8.0, 10 mM NaCl, 0.2% NP-40) followed by nuclear lysis buffer (50 mM Tris-Cl pH 7.5, 10 mM EDTA, 1% SDS).

Chromatin was sheared using an M220 ultrasonicator and immunoprecipitated with antibody-bound Dynabeads. Unbound DNA was removed via sequential washes with low salt (0.1% SDS, 1% Triton X-100, 2 mM EDTA, 20 mM Tris-Cl pH 7.5, 150 mM NaCl), high salt (same buffer with 500 mM NaCl), and LiCl buffer (1% sodium deoxycholate, 1% NP-40, 1 mM EDTA, 10 mM Tris-Cl pH 7.5, 0.25 M LiCl).

Bound DNA was eluted in elution buffer (1% SDS, 100 mM sodium bicarbonate) by overnight incubation at 65°C. DNA was purified following RNase A and Proteinase K treatment using phenol:chloroform:isoamyl alcohol extraction and quantified. ChIP DNA was analyzed by qPCR using primers listed in Supplementary Table 1. Enrichment was calculated as described by Cock-Rada et al. (2012).

### Data Availability

Mass spectrometry proteomics data are available through the MassIVE Consortium using the MassIVE Dataset Submission (1.3.2) workflow under the dataset identifier MSV000097852.

## Acknowledgments

The authors thank the Director, NIAB, for the core grant, essential resources, and continued support. Mr. Shashikant D. Gawai’s expert assistance with Airyscan microscope is much appreciated. We acknowledge fellowship support from DBT, CSIR, and UGC for PhD students (SS, MS, DD, AS, SK, VMA). This study was partially funded by SERB (EMR/2017/001513) and ANRF (CRG/2022/004361) grants awarded to PS.

Author contributions

SS, MS, AS, VB, and PS contributed to writing—review and editing; SS, MS, AS, and PS were responsible for writing the original draft; SS, DD, MS, AS, and PS handled visualization; SS, DD, MS, AS, VB, and SK performed validation; PS provided supervision; SS, DD, MS, AS, VB, SK, and PS contributed to software development; SS, DD, MS, AS, VB, and SK developed the methodology and carried out the investigation; SS, DD, MS, AS, VB, SK, and PS conducted formal analysis and managed data curation; PS was responsible for conceptualization, resource provision, and project administration. The authors acknowledge the use of AI-based tools (e.g., ChatGPT by OpenAI) for assistance in improving the clarity and language of the manuscript. All content was reviewed and validated by the authors.

